# Scalable Fabrication of an Array-Type Fixed-Target Device for Automated Room Temperature X-ray Protein Crystallography

**DOI:** 10.1101/2024.09.30.615838

**Authors:** Sarthak Saha, Yaozu Chen, Silvia Russi, Darya Marchany-Rivera, Aina Cohen, Sarah L. Perry

## Abstract

X-ray crystallography is one of the leading tools to analyze the 3-D structure, and therefore, function of proteins and other biological macromolecules. Traditional methods of mounting individual crystals for X-ray diffraction analysis can be tedious and result in damage to fragile protein crystals. Furthermore, the advent of serial crystallography methods explicitly require the mounting of large numbers of crystals. To address this need, we have developed a device that facilitates the straightforward mounting of protein crystals for diffraction analysis, and that can be easily manufactured at scale. Inspired by grid-style devices that have been reported in the literature, we have developed an X-ray compatible microfluidic device that can be used to trap protein crystals in an array configuration, while also providing excellent optical transparency, a low X-ray background, and compatibility with the robotic sample handling and environmental controls used at synchrotron macromolecular crystallography beamlines. At the Stanford Synchrotron Radiation Lightsource (SSRL), these capabilities allow for fully remote-access data collection at controlled humidity conditions. Furthermore, we have demonstrated continuous manufacturing of these devices via roll-to-roll fabrication to enable cost-effective and efficient large-scale production.

## 2. Introduction

While cryo-electron microscopy^1^ has recently garnered substantial popularity within the structural biology community, X-ray crystallography remains one of the most used techniques.^2–4^ It continues to unveil atomic-level details of biomacromolecules, playing a crucial role in elucidating their functions. By determining the 3-D arrangement of atoms within these molecules, crystallography provides essential insights into the mechanisms underlying biological processes and enables the rational design of pharmaceuticals and other biotechnological applications.^5,6^

Traditionally, crystallographers have relied on laborious procedures to optimize, grow and mount high-quality protein crystals.^7,8^ These crystals are often fragile and sensitive to external factors, such as temperature fluctuations, dehydration, and mechanical stress. Moreover, the exhaustive analysis of easily accessible targets has shifted the focus towards more challenging proteins, such as membrane proteins, where the ability to grow even a small crystal can be a success. Thus, the increasing focus on bioengineering and drug development efforts involving smaller protein crystals has created a need for methods capable of studying them. To analyze small and delicate crystals, both multi-crystal and serial crystallography methods have emerged as valuable techniques.^9,10^ Multi-crystal analysis is performed by stitching together partial datasets from multiple crystals, with the number of crystals depending on the amount and quality of the resulting data. Serial crystallography experiments represent the limit of a multi-crystal approach, with only a single diffraction image collected per crystal, and images from hundreds to thousands of crystals combined to produce a complete dataset. While serial crystallography was first developed for use at X-ray free electron laser (XFEL) sources, it has increasingly been used at synchrotrons as well. In particular, both serial and multi-crystal methods facilitate room temperature (RT) data collection, which can help access the dynamics of protein motion at physiologically relevant temperatures.^10–13^ However, these methods come with the challenge of needing to efficiently deliver numerous crystals to the X-ray source while maintaining their stability at non-cryogenic conditions.

There are two main methods of delivering crystals in a high-throughput fashion for serial crystallography.^14–16^ First is the use of microfluidic injectors that shoot a liquid jet containing a slurry of crystals into the path of the X-ray beam.^14,17–21^ However, this technique can suffer from the problem of low hit rates (one relatively early report estimated one hit per 1600 crystals) when working with slower pulsed X-ray sources,^22^ thus requiring very high sample consumption and limiting its applicability to only those protein crystals that can be produced in large quantities. Some of these injectors use viscous transport medium to combat the hit rate,^21,23–25^ but it is accompanied by intensive sample handling. These hit rate issues become more significant when trying to perform similar injector-based experiments at synchrotrons due to the lower flux of X-rays available, and therefore lower data acquisition rates compared with XFELs. There are also added complexities arising from the setup of liquid jet systems, including the proper alignment, maintenance and cost issues. Given the higher accessibility of synchrotrons compared to XFELs, structural biologists are more likely to take advantage of synchrotrons and a complicated microfluidic injector might not be the most likely option.

Fixed-target mounting strategies are an alternative option that has the potential to dramatically reduce sample consumption.^17,22,26–31^ Protein crystals are mounted in the fixed-target device, preferably in an array form that can be rapidly translated to allow the X-ray beam to optimally target each crystal in turn. Thus, it is possible to achieve a high hit rate and low sample consumption, making multi-crystal and serial data collection at the synchrotron or XFEL more feasible.

However, to minimize the amount of device material interacting with the X-ray beam, a significant number of fixed-target devices are not sealed. The use of such open-faced devices requires careful sample handling and storage to avoid crystal dehydration. Furthermore, such devices require the use of a humidified stream of air to maintain crystal quality during data collection. These humidified setups are gradually being adopted at synchrotrons around the world, coupled with robotic sample handling capabilities that allow for transfer from a humidified storage compartment in a rapid fashion.^32,33^ Unfortunately, the current landscape of fixed-target chip technology is such that most robotic systems are set up only for cryogenic samples and thus devices for RT data collection still require manual loading onto the goniometer at the beamline. However, robotic sample handling systems, like the Stanford Automated Mounter (SAM) used at both the Linac Coherent Light Source (LCLS)^34^ and Stanford Synchrotron Radiation Lightsource (SSRL),^35^ have the potential to overcome this challenge. SAM has been upgraded for RT remote-access exchange of samples at controlled-humidity conditions, which are shipped and stored within SAM-compatible crystallization plates (Figure 5c). The use of SAM is highly efficient as it impacts the throughput of the process eliminating the need to manually swap samples inside the hutch.

As mentioned above, one of the challenges in X-ray sample delivery is reducing background scatter, whether from the device material in fixed-target methods or the liquid stream in injectors. While it is particularly difficult to minimize background scatter from liquid streams due to the complex hydrodynamics at small scales, addressing this issue in fixed-target devices is more feasible. Commonly used materials for fixed targets include polydimethylsiloxane (PDMS) and silicon nitride – both of which contain heavier atoms, such as silicon, that can attenuate X-rays and increase background scattering. Hence, there is a need to move towards either ultra-thin micro-manufactured hard materials or low atomic number materials like plastics. Recent studies have used various materials including PDMS, silicon nitride, polycarbonate, and commercially available MiTeGen polyimide MicroMeshes^TM^.^36–38^ These materials are widely recognized in the scientific community, and a substantial body of literature exists detailing their processing methods; however, they have their share of drawbacks. The crystalline nature of hard materials such as silicon suffers from the potential problem of stray diffraction signals from the mount itself, while polymeric materials that are processed by extrusion tend to have increased scattering due to alignment of the polymer chains from the processing.

Beyond considerations related to the interaction of device materials with X-rays, the microscale features and thicknesses of these devices has required manufacturing strategies that grew out of semiconductor industry. Thus, devices are produced through batch processes involving etching or lithography techniques, resulting in a considerable cost factor for these devices. Additionally, many of the currently available fixed-target devices exhibit suboptimal or negligible optical transparency, resulting in challenges during both loading and data collection. For example, silicon-based devices are opaque, rendering optical mapping of crystal locations infeasible. Consequently, data collection must be conducted at every potential crystal location in the device, a process that is both time-consuming and inefficient, as not all sites contain crystals. An alternative approach is rastering, which involves scanning the sample with low-dose X-rays across all positions. However, this method also has its limitations, as it requires exposing each position to radiation, albeit at a lower dose. While some materials, such as polyimide-based devices offer improved optical transparency, the inherent orange tint restricts the feasibility of conducting simultaneous spectroscopic analyses. To circumvent these issues, we have utilized an alternate class of polymeric materials that offer enhanced cost-effectiveness, simplified manufacturing processes, superior optical properties, reduced signal attenuation, and high X-ray transparency.

Here, we report the design, construction, and performance evaluation of our novel array-type fixed-target device for X-ray crystallography (AFD-X). We offer an in-depth exploration of the new polymeric materials with superior optical properties, X-ray compatibility, and low-cost roll-to-roll fabrication techniques that can be used to manufacture a large number of devices. We also describe practical guidelines for the utilization of the AFD-X. We have showcased a streamlined process of protein structure determination, starting from sample loading, shipping crystals on-chip from the crystallization laboratory to the synchrotron facility, and automated remote data collection at room temperature using seamless integration with the current robotic pipeline at SSRL. We have validated our approach with *Gallus gallus* hen egg white lysozyme, *Thaumatococcus daniellii* thaumatin and *Tritirachium album* proteinase K.

## 3. Materials and Methods

### 3.1. Photomask Design

Our design required multilayered photolithography either to fabricate the SU-8 based devices or the moulds for NOA based devices, hence photomasks were designed for the two layers in Adobe Illustrator and were made to fit on a 3” silicon wafer (provided in SI). Each photomask included fiducial markers that allowed for alignment during multilayer photolithography. Each design was printed on a polyester film on a right-reading down manner (Fineline Imaging, Colorado Springs, CO, USA) at 10,160 dpi.

### 3.2. Batch Device Fabrication

We used two different types of batch fabrication methods. The first of these involved sequential patterning of a UV-sensitive polymer, similar to methods described in our previous papers.^39,40^ The second method allowed for direct moulding polymer in a single step using a different UV-sensitive polymer. This approach allowed for small scale validation of our methodology before transitioning to continuous roll-to-roll manufacturing.

#### 3.2.1. Batch Fabrication Using SU-8

Multilayer photolithography was performed using two grades of negative-tone photoresist (SU-8 2100 and SU-8 2002) and a sacrificial lift-off layer (LOR 3A), purchased from Kayaku Advanced Materials (formerly Microchem, Westborough, MA, USA) and processed using the standard manufacturer’s guidelines. Propylene glycol methyl ether acetate (PGMEA), used for the development of SU-8, was purchased from Sigma-Aldrich. RD6 developer (tetramethylammonium hydroxide) for LOR 3A was purchased from Futurrex (Franklin, NJ, USA). Silicon wafers (76.2 mm, Type P, test grade, 100 orientation) were purchased from University Wafer, Boston, MA, USA.

Firstly, silicon wafers were cleaned with nitrogen gas to remove particulates and/or dust from the surface. The silicon wafer was first coated (Laurell WS-650-23) with a sacrificial layer of 100 nm LOR 3A at a spin speed of 3000 rpm for 60 s, followed by baking at 185°C for 5 min. Next, a 2 µm layer of SU-8 2002 was deposited by spinning at 3000 rpm for 60 s. This layer acted as the base of the device and contains holes to wick away excess fluid (Figure 1). This 2 µm layer was prebaked for 1 min at 95°C and patterned using the photomask containing designs for the “wicking hole” using standard photolithography. The mask aligner (SUSS MA/BA 6, UV lamp wavelength – 365 nm, power 9.6 mW/cm^2^) was set for an 8 s exposure to achieve a total exposure dose of 80 mJ/cm^2^. This was followed by a 1 min post exposure bake at 95°C. Next, we spin coated a 100 µm layer of SU-8 2100 at 3000 rpm. This was followed by a two-step prebake starting at 65°C for 5 min followed by 95°C for 20 min. The second photomask, containing the design for the “crystal wells,” was aligned to the first patterned layer using fiducial makers and was exposed for 24 s to achieve a total exposure dose of 240 mJ/cm^2^. Following this, the sample was post-baked at 65°C for 5 min and 95°C for 10 min. After allowing the wafer to cool for 5 min, the two layers of SU-8 were developed in PGMEA to dissolve the uncured SU-8, revealing the crystal traps and wicking holes. Finally, the wafer was dried with nitrogen and developed overnight in RD6 to wash away the sacrificial LOR layer, resulting in a free-standing SU-8 structure, which was air dried.

**Figure 1.**
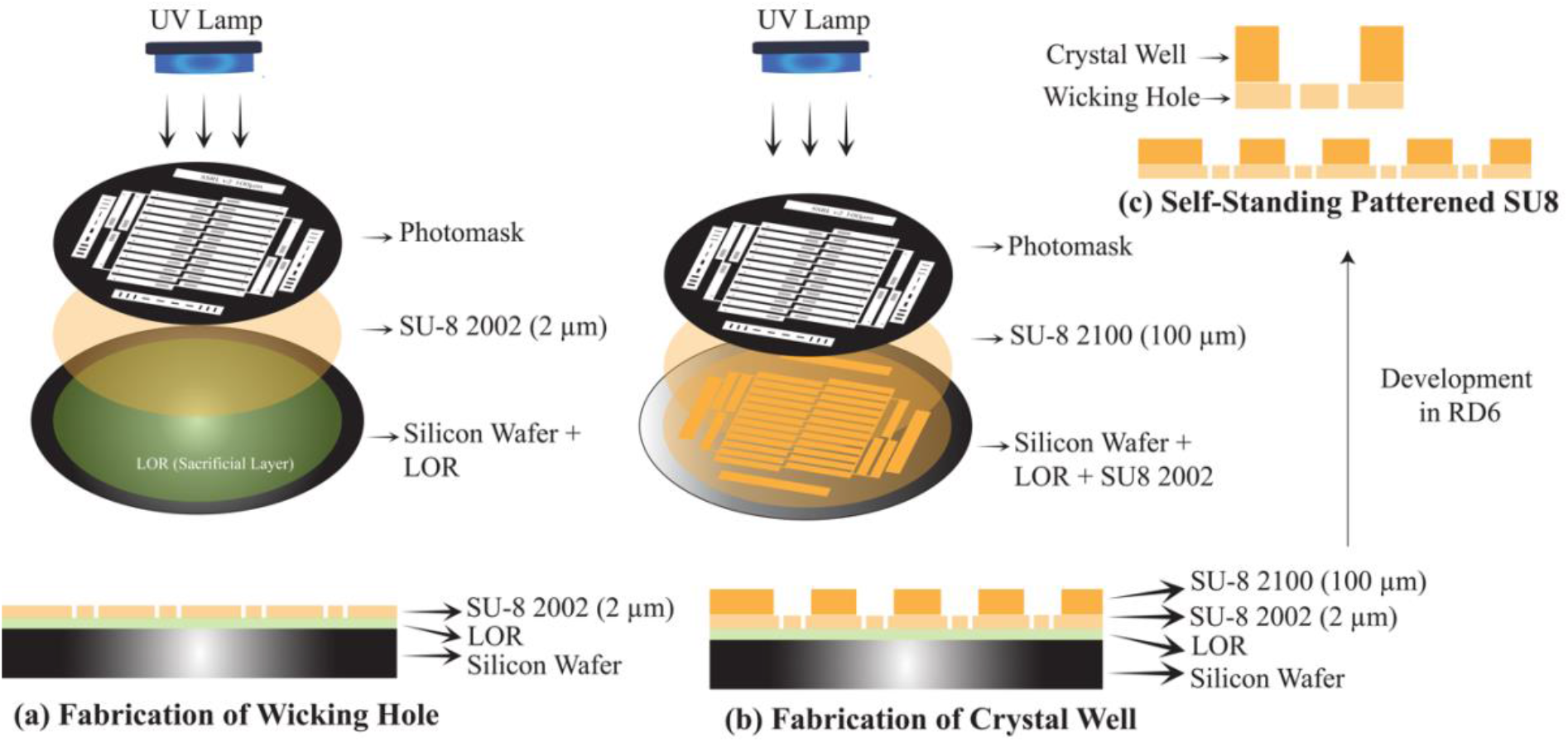
Schematic showing multilayer fabrication of SU-8 based grids on a silicon wafer using photolithography. **(a)** First, the wicking hole layer is fabricated by using 2 µm SU-8 2002 followed by **(b)** the crystal trap layer using 100 µm SU-8 2100. **(c)** Finally, the fabricated device is released from silicon wafer to form self-standing patterned SU-8 grids.

#### 3.2.2. Batch Fabrication Using NOA

Nanoimprinting was performed using NOA 68T (Norland Products, Jamesburg, NJ, USA) with a polydimethylsiloxane (PDMS) (Ellsworth Adhesives, Germantown, WI, USA) master. The flexibility of the PDMS allows for easy separation of the device from the mold after curing. To create the PDMS master, multilayer photolithography was first used to create an inverse SU-8 on-silicon master using photolithography (Figure 2a). Here, similar procedures were used as above, except that the sacrificial layer of LOR 3A was not included since we wanted to retain the SU-8 layer on the wafer surface. Additionally, a 25 µm thick layer of SU-8 2025 was used to create the mold for the wicking holes, rather than the 2 µm thick layer used for direct photolithography, because when the pattern was transferred onto NOA, a self-supporting structure of NOA could not be achieved using 2 µm thick features due to its insufficient strength. This was followed by 100 µm of SU-8 2100 and patterning with the “crystal well” photomask. Baking and exposure time were followed as per the manufacturer’s guidelines, as above. The silicon wafer containing SU-8 master was silanized using 150 µL (tridecafluoro-1,1,2,2-tetrahydrooctyl)trichlorosilane (Gelest Inc., Morrisville, PA, USA) via vapor treatment. 150 µL silane and the SU-8 master were placed in continuous vacuum for 5 minutes, followed by closing of the valve to preserve vacuum in the closed desiccator overnight.

**Figure 2.**
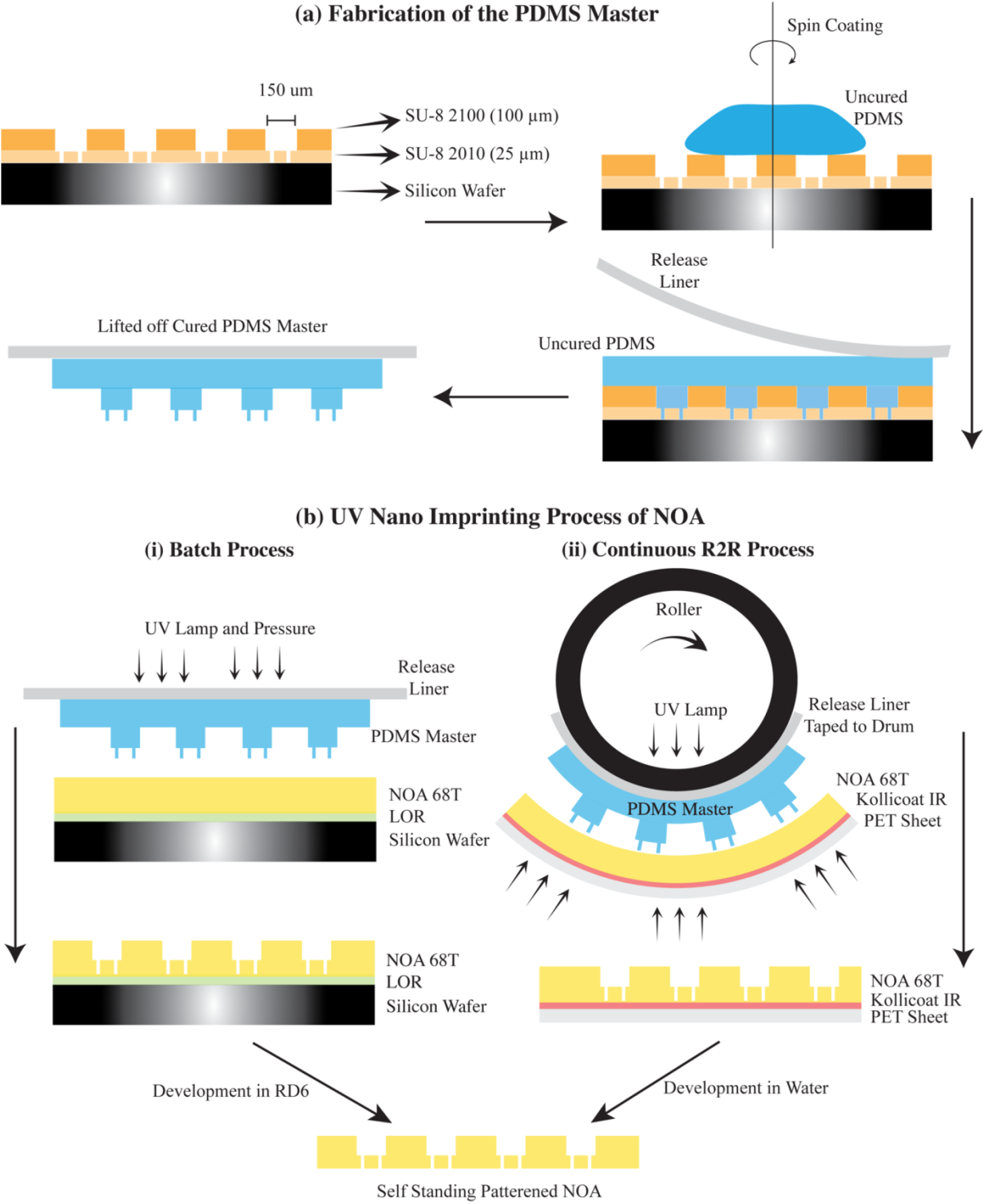
Schematic showing the fabrication of grids using NOA. **(a)** negative PDMS master is fabricated by micro molding off SU-8. **(b)** The PDMS master is used to imprint the design into NOA 68T using both **(i)** batch and **(ii)** continuous roll-to-roll processing.

To make the PDMS mould, PDMS (SYLGARD^TM^ 184 Silicone Elastomer) was mixed with the curing agent in a 10:1 mass ratio and degassed for 1 hour. This was spin coated on the SU-8 master at 500 rpm for 60 s followed by curing at 90°C for 1 hour. Next, another layer of PDMS was coated at 500 rpm. The uncured layer was carefully covered with a silicone release liner C1S (MPI Release, Winchester, MA, USA) and left undisturbed for 48 hours to allow the PDMS to cure at room temperature. The release liner containing the PDMS mould was then peeled off of the SU-8 master and was ready for nanoimprinting. The thin PDMS master was required for compatibility with roll-to-roll processing, though a thicker master could be used for simple batch processing.

As shown in Figure 2, to fabricate the device out of NOA 68T, a fresh silicon wafer was coated with a sacrificial layer of LOR3A at 3000 rpm followed by baking at 185°C for 5 min. Next a layer of NOA 68T (Norland Products Inc., Jamesburg, NJ, USA) was coated at 3000 rpm resulting in 90 µm thickness. The coated wafer was then covered with the PDMS mould and inserted into the nanoimprinter (Nanonex, Monmouth Junction, NJ, USA). Nanoimprinting uses a combination of pressure, temperature and UV to transfer the pattern from master to substrate, the conditions used were 25°C, 2 psi (13.79 kPa) and 1 min of UV exposure. Following transfer of the pattern into the NOA, the silicon wafer was immersed in RD6 overnight to remove the LOR layer, releasing the NOA device.

### 3.3. Continuous Roll-to-Roll Device Fabrication

NOA has an added advantage of being compatible with roll-to-roll (R2R) processing, which is helpful for larger-scale continuous manufacturing. Stamps for R2R imprinting were prepared using PDMS and a C1S release liner as described above for batch nanoimprint lithography. A R2R Nano Emboss 200 (Carpe Diem Technologies, MA) was used for UV-assisted roll-to-roll nanoimprinting. The fabricated stamp was attached to the 15.6 cm quartz drum which surrounds a 365 nm UV LED lamp. Optical grade 125 µm PET film was coated with a sacrificial layer of Kollicoat® IR (Sigma-Aldrich) using a R2R Nano Emboss 200 (Carpe Diem Technologies, MA). This coated PET film was used as the substrate material. NOA 68T was then coated onto the PET using a Meyer rod attachment on the Nano Emboss 200 instrument. 365 nm wavelength UV light with 2.8 W/in^2^ power density was used to cure the NOA. The imprinting line was run at 12 in/min (300 mm/min) without any additional pressure. The tension inherent in the R2R machine itself was sufficient to achieve the desired pressure for effective imprinting. Finally, the PET sheet was placed in a water bath overnight to dissolve the sacrificial Kollicoat® IR layer and release the self-standing NOA layer.

### 3.4. Device Assembly and Mounting

The molded devices were attached to the magnetic base portion of a copper-magnetic mount (Crystal Positioning Systems, Jamestown, NY, USA) using a support structure made from single-sided adhesive tape (Adhesive Research® 7759). This tape is a 127 µm-thick polyester film coated with 30 µm thick acrylic adhesive. The tape was cut into 2.3 mm × 14 mm pieces using a plotter cutter (Graphtec CE6000-40). The crystal trap side of the NOA and SU-8 devices was manually bonded to the adhesive side of the tape. The assembled chip, as shown in Figure 3a, was then glued to the magnetic pin base using universal puck base plate assembly and a 3-D printed jig to facilitate alignment (Figure 3c). The 3-D printed assembling jig was designed in Solidworks (attached in SI) and printed using EOS P110 printer and nylon-11 (PA11 white). Upon completing assembly, a black spot was placed on the top side of the mount, to help with easy visualization of device orientation.

**Figure 3.**
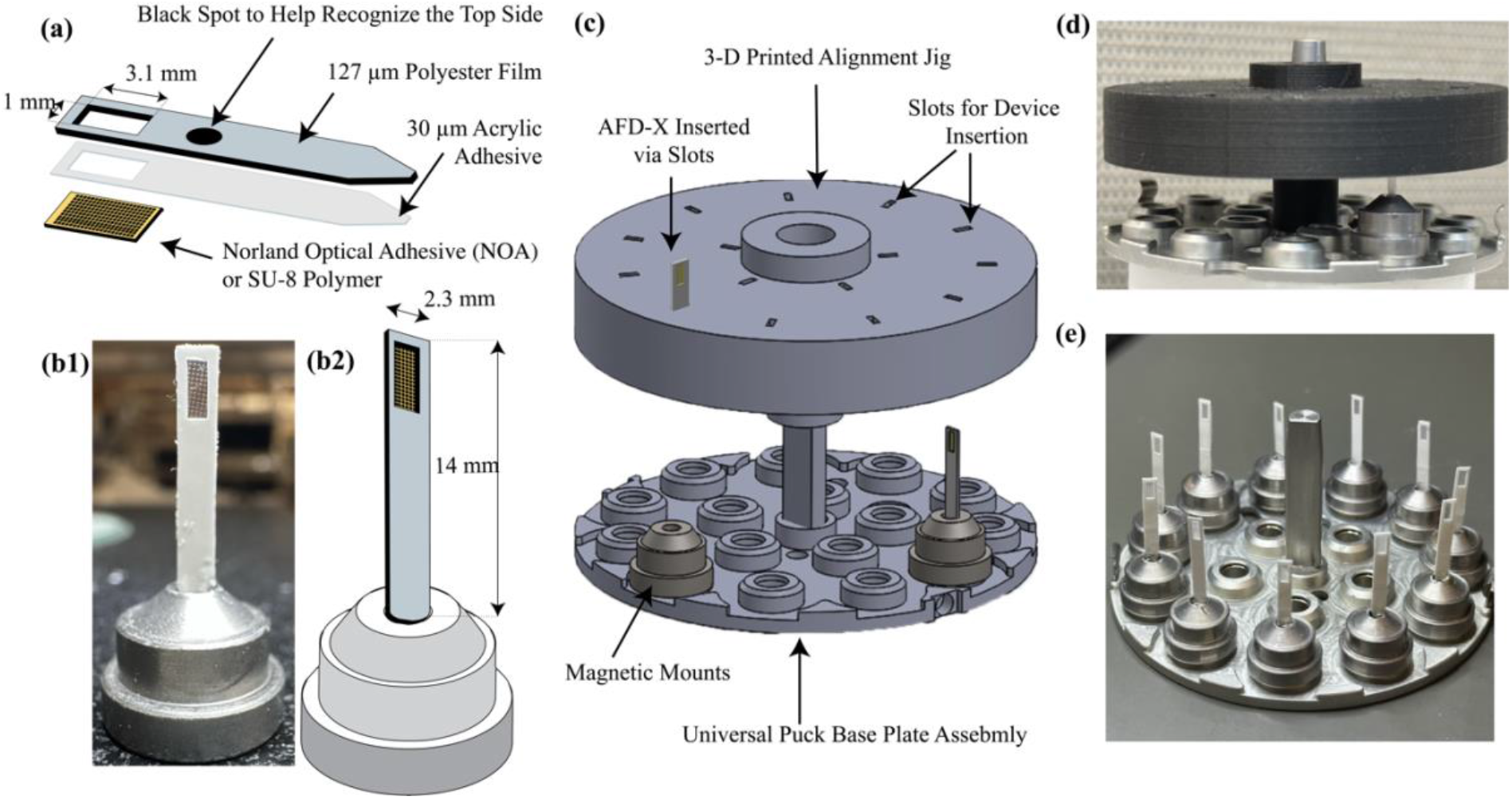
**(a)** Schematic showing the array-type fixed-target device for X-ray crystallography (AFD-X) with the X-ray transparent polymer sample array attached to the single-sided adhesive tape. **(b)** Photograph and schematic of the AFD-X glued to a magnetic mount. **(c)** 3-D model of the jig used to assemble the device **(a)** onto the magnetic mount. **(d)** Photograph of the assembly jig in use. **(e)** A batch of final devices ready to be loaded with protein crystals.

### 3.5. Device Characterization

Optical micrographs of the device were captured using a Microscope Illuminator 385B (Bausch & Lomb Model – ASZ45L3) and a Zeiss SteREO Discovery V12 stereomicroscope. The height of the various device features was measured using a Nexview 3-D Optical Profilometer (Zygo Corporation, CT, USA).

### 3.6. Protein Crystallization

#### 3.6.1. Lysozyme Crystallization

*Gallus gallus* hen egg white lysozyme was purchased from Hampton Research Inc. and was dissolved in 50 mM sodium acetate buffer (Fisher Scientific, ACS grade) at 200 mg/mL, at pH 4.6. A precipitant solution of 30% w/v polyethylene glycol monomethyl ether 5,000 (PEG MME), 1.0 M sodium chloride, 0.05 M sodium acetate trihydrate pH 4.6 was purchased from Hampton Research. Crystallization trials were set up at ambient laboratory temperature (∼23°C) using a sitting drop strategy using Cryschem plates (Hampton Research Corp.) with 2 µL protein and precipitant solution each and 1 mL of reservoir solution.

#### 3.6.2. Thaumatin Crystallization

Thaumatin from *Thaumatococcus daniellii* (T7638) and potassium sodium tartrate were purchased from Sigma Aldrich. N-(2-acetamido)iminodiacetic acid (ADA buffer) was purchased from Fisher Scientific. Thaumatin in DI water at a concentration of 25 mg/mL was used as the protein solution. 1 M potassium sodium tartrate in 0.1 M ADA buffer at pH 6.5 was used as the precipitant solution.^41^ Crystallization trials were set up in a similar manner to the lysozyme crystallization protocol.

#### 3.6.3. Poteinase K Crystallization

Proteinase K from *Tritirachium album* (P2308), ammonium sulphate, Tris-hydrochloride (Tris-HCl), and 4-(2-hydroxyethyl)-1-piperazineethanesulfonic acid) (HEPES buffer) were purchased from Sigma Aldrich. 20 mg/mL of protein was mixed in 50 mM HEPES at pH 7.1.2 M ammonium sulphate in 0.1 M Tris-HCl at pH 8 was used as the precipitant solution.^41^ Crystallization trials were set up similar to lysozyme.

### 3.7. Device Operation and Data Collection

SSRL crystallization plates (Crystal Positioning Systems, Jamestown, NY, USA) were prepared by placing the liner cup and foam insert in each well. 300 µL of reservoir solution was pipetted into the foam insert and was sealed using Crystal Clear tape, allowing the chamber to equilibrate for 10 minutes. 0.5 µL of protein crystal slurry was pipetted out of the sitting drop and deposited into the grids of the AFD-X. The excess liquid was wicked through using a Kimwipe from the bottom of the AFD-X (as shown in supplementary video), leaving the crystals in the grid of AFD-X (as shown in Figure 5b). Next, the AFD-X containing crystals was placed inside the equilibrated SSRL crystallization plate and sealed again with Crystal Clear Tape. As shown in Figure S3, the crystallization plates were loaded into the SSRL Thermal Shipping Kit (Crystal Positioning Systems, Jamestown, NY, USA), which reports temperature control of the samples for up to 168 hours. AFD-X containing crystals were shipped via overnight courier service to SSRL. Crystals at room temperature were transferred from inside SSRL crystallization plates onto the beam line goniometer using the Stanford Automated Mounter (SAM).^35^ Data collection was performed at beamline 12-1 using an X-ray wavelength of 0.98 Å, exposure time of 0.2 s and a beam size of 40 × 50 µm using a transmission factor of 1%. Crystals were maintained at room temperature (23°C) and a set humidity level of 96% using an Arinax humidity control nozzle (Yongin-si, Gyeonggi-do, South Korea). The AFD-X assembly was mounted perpendicular to the X-ray beam (defined as 90°), and data was collected from 30° to 150°, resulting in 120° of data per crystal using 0.5° of oscillation per frame. An Eiger2 XE Pixel Array Detector was used at a sample-to-detector distance of 160 mm.

### 3.8. Data Processing

Diffraction data sets were integrated, merged, and scaled using HKL-3000.^42^ The structures were solved by molecular replacement (MR) with Phaser^43^ using the coordinates of 8f00 as a search model for lysozyme, 5×9m as the model for thaumatin and 8f07 for proteinase K.^41^ The data refinement was performed with REFMAC5^44,45^ and structures were built into electron density using Coot.^46^ Nika software for 2-D diffraction data reduction in Igor Pro (Wavemetrics Inc.) was used to analyze the background scattering.^47,48^ Integration was performed in 2θ with log binning using the calibrated beam center and sample-to-detector distance.

## 4. Results and Discussion

The primary objective of this study was the development of a user-friendly array-type fixed-target device that can be produced with ease in a cost-effective manner. Furthermore, we sought to employ more user-friendly materials compared to those utilized in previous endeavors. In pursuit of this goal, we have showcased the applicability of R2R manufacturing techniques alongside photolithography using polymeric materials. One of the polymers: SU-8, can be readily fabricated using photolithography via batch manufacturing strategies. The second material, NOA has the potential to be manufactured through nanoimprinting, both in batch and continuous R2R processing. These polymeric materials were chosen based on their superior optical clarity, low X-ray background, and UV curable properties, which further enhances their suitability for integration into fixed-target devices.

Our overarching aim is to streamline the X-ray crystallography workflow, commencing from the cultivation of protein crystals within laboratory settings and culminating in the collection of room temperature data at X-ray facilities, all accomplished in a cost-efficient and user-friendly manner. To realize this objective, we set forth specific design criteria to ensure compatibility with: (i) SSRL crystallization plates, (ii) the humidity control systems at SSRL, and (iii) the robotic arm used for room temperature data collection. Although the primary design focus was on room temperature data collection, it is noteworthy that AFD-X are also adaptable for cryogenic data collection.

### 4.1. Device Design and Batch Processing

In the context of designing X-ray-compatible sample holders, our prior works^39,40^ have demonstrated the suitability of SU-8, a common UV-curable photoresist used in microfluidic device manufacture. However, SU-8 is inherently constrained by its limited feasibility for continuous R2R manufacturing due to the extensive thermal baking processes required before and after UV exposure, thus increasing the manufacturing costs and time expenditures.

In light of these limitations, we aimed to identify an alternative UV-curable polymer that would obviate the need for thermal baking steps. In our pursuit, we identified Norland Optical Adhesives (NOAs), as a promising class of materials that aligned with our requirements. Among the NOA variants, NOA 68T emerged as particularly well-suited for our application due to a viscosity that was optimized for the range of thicknesses (50-150 µm) relevant to our device designs. NOA polymers can be readily cured through simple UV exposure, resulting in crosslinking that makes them amenable for R2R use.^49^

An important design criterion for developing grid-based devices is the need for thin materials in the area that acts as the X-ray window to minimize scatter and signal attenuation. When fabricating the grids out of SU-8, we employed a 2 µm-thick X-ray window. However, when making the device out of NOA, it needed a thicker 20 µm-thick layer of NOA 68T, due to the weaker mechanical properties of cured NOA 68T, as compared to SU-8. Specifically, when attempting to create a 2 µm-thick structure for a 30 × 30 µm X-ray window, NOA 68T exhibited structural instability and disintegration.

From a processing perspective, to create through-holes in a 20 µm-thick layer, it proved necessary to create a negative PDMS master with taller, 25 µm features (Figure 4). This additional 5 µm height allowance was required for effectively excluding the NOA 68T material from the bottom of the features, thereby facilitating the creation of through-holes when using a 2 psi pressure for processing. Both designs also included 150 µm crystal traps featuring 30 µm wicking holes (as shown in Figure 4) to allow for efficient removal of excess mother liquor. The height of the resulting features was obtained using optical profilometery, which allowed us to optimize parameters such as PDMS master thickness, nanoimprinting pressure as well the grade to NOA used in batch fabrication prior to moving on to roll-to-roll manufacturing.

**Figure 4.**
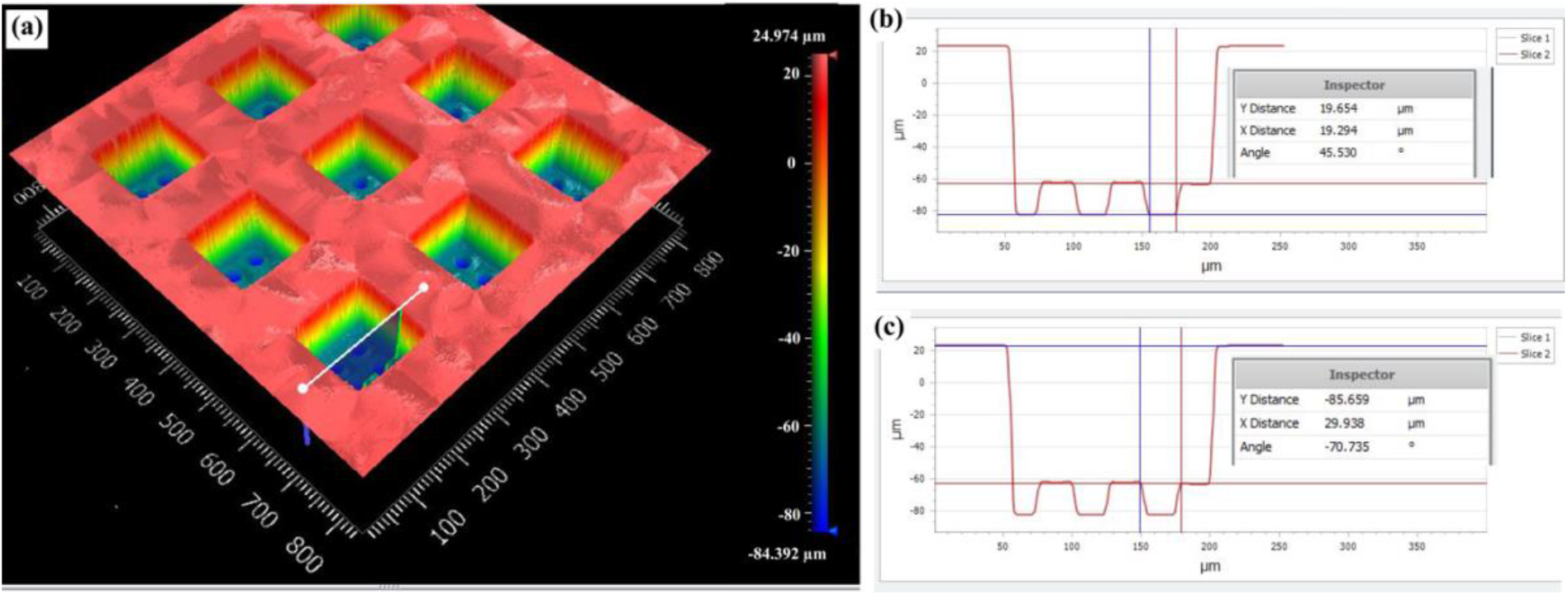
**(a)** 3D Profilometry image of grids corresponding to AFD-X chip. **(b-c)** Line trace depicting the 105 µm thickness of the open faced AFD-X along with the 150 µm-wide crystal traps and 30 µm-wide wicking holes.

### 4.2. Roll-to-Roll Processing and Lower Associated Manufacturing Cost

R2R processing was achieved using PET as a substrate and NOA as the polymeric material. The NOA was continuously coated onto the PET and then passed through a roller that contained the PDMS master with the negative mould. The UV light embedded within the roller then simultaneously cured and moulded the NOA. The feature sizes in NOA were dictated by thickness of the coated NOA and size of the PDMS mold. To optimize R2R manufacturing, we conducted systematic experiments focusing on key variables such as the applied pressure during imprinting, the intensity of UV exposure, and the dimensions of the PDMS master. Our objective was to attain the highest possible precision in the layer thickness during the R2R process. At a constant roll speed of 30 cm/min, we determined that an ideal UV intensity of 0.434 W/cm^2^ was necessary for successful imprinting. Below this threshold, we experienced incomplete curing of the NOA.

Regarding the application of pressure during the imprinting process, we observed that the tension inherent in the R2R machine was found to be sufficient to achieve effective imprinting. Interestingly, any additional pressure applied to the drum where the PDMS master was affixed resulted in buckling and/or deformation of the pillars used to create the through-holes (Figure S1b). This result is a consequence of the flexible and elastomeric nature of the PDMS master, and the use of alternative materials for the master would necessitate a re-evaluation of processing conditions.

Similar to batch fabrication, it proved necessary to make the wicking hole features on the PDMS master taller than their intended depth to ensure that the holes would be open. While the mechanical properties of NOA necessitated the same 20 µm thick layer for the wicking hole layer, a different issue emerged if we increased the PDMS feature height to 50 µm. In this case, some of the pillars within the PDMS master exhibited bending behavior (similar to the deformations that were observed upon the application of pressure in batch, shown in Figure S1b), resulting in anomalies within the imprinted structures. This observation highlights the delicate balance required when determining the dimensions of various features to facilitate efficient pattern transfer during the nano-imprinting process.

Employing R2R technologies significantly expedited the manufacturing process compared to batch processing. For instance, with R2R processing, we achieved imprinting at a rate of 300 mm/min. By employing four PDMS masters attached to a 300 mm diameter roller, we could produce nearly one full rotation worth of devices in 3 minutes, resulting in 42 × 42 mm worth of imprinted structures, or approximately ∼50 of our individual AFD-X. Consequently, within 60 minutes, we could manufacture approximately 80 batches corresponding to 4000 AFD-X. In contrast, while using a batch process, we would only be able to produce a single SU-8-based batch or four NOA-based batches of AFD-X in the same period of time. It should be noted that these timeframes do not include the overnight duration typically required for the lift-off process.

The cost benefits of transitioning to continuous R2R manufacturing are substantial as well. Aside from the fixed cost of polymer materials, the primary difference in operational costs lies in the raw material—specifically between the silicon wafers used in batch processing and the PET sheet used in R2R. In 60 minutes of R2R operation, approximately 3.60 meters of PET sheet is used, costing $2.25. This results in a cost per batch of $0.028 ($0.00056 per AFD-X). In contrast, a single silicon wafer used in batch processing costs $10 and can be reused only 4-5 times, bringing the cost per batch to $2 ($0.04 per AFD-X). Thus, the R2R process is roughly 70 times cheaper in terms of raw material costs. Additionally, the efficiency gain is significant in terms of labor. The R2R process produces 80 batches in one hour compared to only 1 batch in the batch process, reducing the labor costs by 80 times. Our approach offers advantages in both economic and manufacturing efficiency aspects.

### 4.3. Device Assembly and Operational Compatibility

After creating the polymer grids, the next critical phase involved ensuring the compatibility of the AFD-X with both the SAM robot and crystallization plate. The SAM robot can handle objects similar in size to the standard pin mounts used in cryocrystallography (around 2.7 × 18 mm). Therefore, the AFD-X was designed to fit these specifications, with a final size of 2.3 × 14 mm to ensure adequate tolerances (as shown in Figure 3b). Upon testing with the SSRL robot, the chips exhibited seamless handling without any issues. The same design features also allowed for the chips to be compatible to the humidity stream which stabilized the protein crystal during data collection, and with cryogenic sample handling and data collection, if needed. The imprinted polymer was attached to single-sided adhesive tape having a window of 1 × 3.1 mm window to provide support since the thin polymer device itself does not have the mechanical strength to remain rigid in the presence of the air flow from the humidity stream. Once the single-sided tapes were prepared and attached with the polymer grids (as depicted in Figure 3a), the subsequent crucial step involved precise assembly with the magnetic mount. Careful alignment was paramount to avoid collisions during handling by the SSRL robot.

### 4.4. AFD-X Loading and Operation

After successfully assembling the AFD-X with the crystal mount, the devices were ready for use. Crystals were cultivated via sitting drop (Figure 5a, S2) and transferred to the grid via pipetting, which allowed for the easy transfer of many crystals at once, thereby eliminating the need for handling of individual crystals. Furthermore, the quick transfer of crystals surrounded by mother liquor allowed the process to be performed under a simple microscope without the need of a humidified environment. Finally, the excess mother liquor was removed by wicking with a Kimwipe (as shown in supplementary video). The simplicity of these handling steps is in contrast to similar array chips made from silicon and/or silicon nitride, which require the use of a specialized vacuum pump setup to wick through liquid due to the lower wetting properties of such chips.^50^

**Figure 5.**
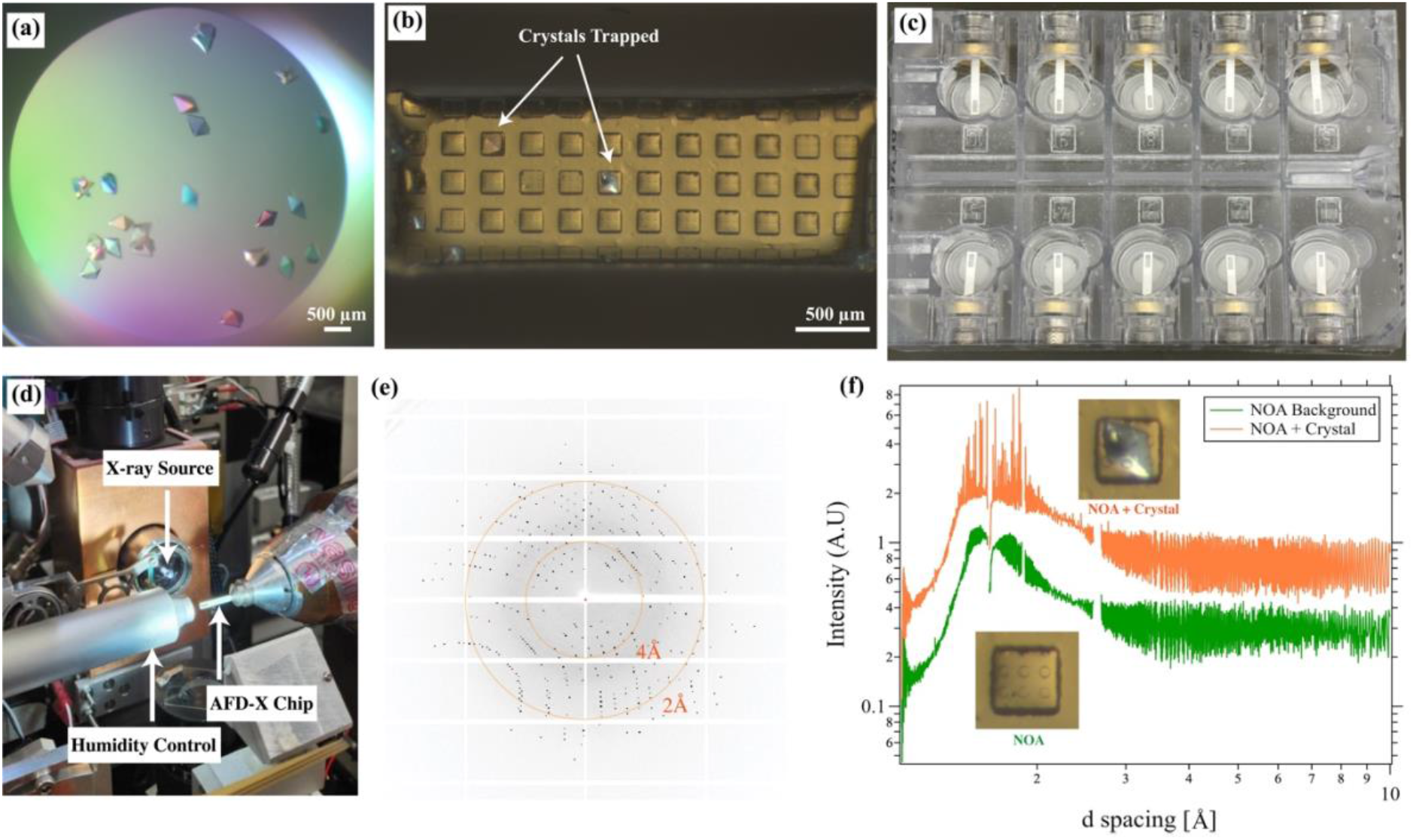
**(a)** Proteinase K crystal grown in a sitting drop and **(b)** transferred to AFD-X chip. **(c)** Photograph of AFD-X loaded with protein crystals inside the SSRL crystallization plate for safe transport. **(d)** AFD-X loaded onto SSRL beamline 12-1 using the robot. **(e)** The 2-D diffraction signal for the lysozyme crystal collected in our AFD-X chip at RT, corresponding to the orange curve in **(f). (f)** Graph of the 1-D integrated X-ray intensity profiles comparing the relative strength of the observed diffraction signal from a lysozyme crystal compared to the noise from background scattering due to the presence of device materials and crystallization solution as a function of resolution.

In addition to easy wicking, the optical transparency of the AFD-X meant that it was possible to directly observe crystals during loading (Figure 5b, S2d) and pipette additional crystals if necessary. This again is in contrast to silicon-based array chips where visualization can only be done in reflection. While such imaging can be done during chip loading, reflection setups are less common at X-ray beamlines, meaning that visualization of crystals during data collection is more challenging, and methods such as low-dose X-ray rastering must be used instead. The transparency of our AFD-X chips enabled selective data collection only at the locations where crystals were present (Figure 5b, S2d). We would also note that this optical transparency has the potential to enable simultaneous spectroscopic studies and/or light-triggered time-resolved studies, though such efforts were beyond the scope of the current work.

### 4.5. Remote and Automated Room Temperature Data Collection

We were able to successfully ship our AFD-X chips containing crystals using SSRL plates and the associated thermal shipper. Since the AFD-X chips are not sealed, it was essential to maintain the samples in a humidified environment to preserve crystal quality during storage, shipping, and data collection. Within the SSRL plate, absorbent material containing 300 µL of precipitant solution served to maintain the crystals during storage and transport. During data collection, a stream of humidified air was used to stabilize the crystals. The relative humidity needed to stabilize a given crystal sample can be considered with regards to the precipitant solution used during crystallization. For each of our three protein targets, the relative humidity was maintained at 96% during data collection. Our studies required 30 minutes per chip to collect data sets from multiple crystals, and during this period we did not visually observe any evidence of dehydration of the crystals owing to the humidity control. Furthermore, the data obtained from multiple crystals on the same chip consistently yielded similar unit cell parameters (Figure S5), reinforcing the stability of the crystals throughout the data collection process.

Although we did not pursue a fully automated and high-throughput data analysis strategy, we would note that the ability to use the SSRL robot meant that the time needed to enter the hutch and manually exchange devices—typically taking 4-5 minutes per change—was eliminated. Leveraging the robot system enabled a swift chip exchange within 1 minute per chip, significantly expediting the process. Furthermore, and the ability to do automated sample exchange means that we were able to perform RT data collection remotely.

### 4.6. Low Background Scattering of AFD-X Chips

To quantify background scattering we measured the diffraction signal in our AFD-X chip. We compared the scattering of AFD-X with existing and commercially available MiTeGen MicroMeshes^TM^, silicon nitride, and polycarbonate chips.^36–38^ We observed that both SU-8 and NOA-based AFD-X chips performed similarly to their polymer counterparts (MiTeGen MicroMeshes^TM^ and polycarbonate chips) with maximum intensity of around 0.8-0.9 A.U. at a resolution of 4.5 Å, and observable variations consistent with differences in material thickness (Figure S4). Next, we compared the scattering of lysozyme crystals in the AFD-X chips with the empty chips (Figure 5f). Here we observed a slight increase in scattering, as would be expected due to the presence of the crystal, with an intensity just above 1 A.U. Importantly, we would note that a comparison of both 2-D diffraction images (Figure 5e) and the corresponding 1-D integrations (Figure 5f) highlights the excellent signal-to-noise levels present when comparing the observed diffraction peaks from the crystal with the diffuse background scattering.

### 4.7. Crystal Stability and Structure Determination

To demonstrate the efficacy of our device we collected data from 7 different crystals each of lysozyme, thaumatin and proteinase K across 15 AFD-X shipped in the SSRL crystallization plates and handled by the SSRL robot. Each of the data sets was indexed and we performed a statistical analysis of the unit cell parameters of all 21 individual crystals which were shipped across the country. We observed highly reproducible unit cell parameters amongst the crystals, which can be credited to the sample stability of the AFD-X chip, as well as the easier sample handling that it enables. The statistical analysis was performed after indexing and geometric refinement using methods described by Liu *et al*.^51,52^ We observed that the coefficient variation for the unit cell parameters *a b*, and *c* was quite low: 0.096%, 0.086% and 0.120%, respectively for lysozyme, 0.028%, 0.044% and 0.038%, respectively for thaumatin and 0.020%, 0.024% and 0.031%, respectively for proteinase K (Figure S4a-c), demonstrating highs level of isomorphism. The variation of unit cell parameters was also calculated using standard Euclidean distance, ▽_j,k_ and we observed that all indexable crystals had a ▽_j,k_ value below 1.6, which is significantly below the cutoff of 3.0 recommended by Liu *et al*. (Figure S4d).^51,52^ For all three protein targets we obtained high resolution (∼1.3 Å) room temperature data with nearly 99% completeness from single crystals (as shown in Table 4.1 and Figure 6) and the data was similar to literature reports.^41,53^ While the geometry of the AFD-X meant that we were limited to collecting 220° of usable data, the high symmetry of our crystals, meant that acquisition of only the initial 110° of data was sufficient to achieve completeness.

**Table 4.1.** Crystallographic statistics for data obtained using the AFD-X at room temperature for lysozyme, thaumatin and proteinase K.

|  | Lysozyme<br>(single crystal) | Thaumatin<br>(single crystal) | Proteinase K<br>(single crystal) |
| --- | --- | --- | --- |
| **Data collection** |  |  |  |
| Space group | P4<sub>3</sub>2<sub>1</sub>2 | P4<sub>1</sub>2<sub>1</sub>2 | P4<sub>3</sub>2<sub>1</sub>2 |
| Cell dimensions |  |  |  |
| $a, b, c$ (Å) | 78.96, 78.96, 38.26 | 58.527, 58.527, 151.652 | 68.40, 68.40, 108.10 |
| $\alpha, \beta, \gamma$ (°) | 90, 90, 90 | 90, 90, 90 | 90, 90, 90 |
| Resolution (Å) | 50 – 1.29 | 50 – 1.37 | 50 – 1.39 |
| $R_{\text{merge}}$ | 0.109 | 0.100 | 0.140 |
| Mean $I/\sigma I$ | 14.9 | 10.9 | 9.0 |
| Completeness (%) | 99.3 | 98.9 | 99.0 |
| $CC_{1/2}$ | 0.985 | 0.986 | 0.945 |
| $CC^*$ | 0.996 | 0.996 | 0.986 |
| **Refinement** |  |  |  |
| Resolution (Å) | 31.56 – 1.29 | 32.02 – 1.37 | 32.61 – 1.39 |
| Unique reflections | 30806 | 55649 | 51711 |
| $R_{\text{work}} / R_{\text{free}}$ (%) | 21.73 / 23.69 | 21.83 / 23.44 | 19.98 / 22.54 |
| Ramachadran plot |  |  |  |
| In preferred regions (%) | 99.21 | 99.02 | 96.75 |
| Outliers (%) | 0.00 | 0.00 | 0.00 |
| R.M.S. deviation |  |  |  |
| Bond lengths (Å) | 0.0058 | 0.0068 | 0.0054 |
| Bond angles (°) | 0.895 | 1.029 | 0.842 |

**Figure 6.**
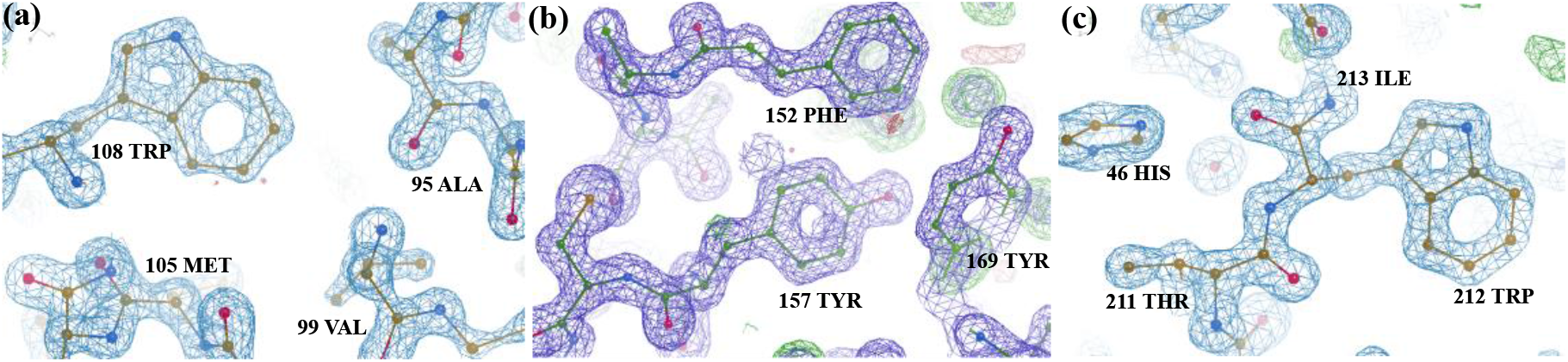
2Fo − Fc electron density map of **(a)** lysozyme, **(b)** thaumatin, and **(c)** proteinase K grown via sitting drop and analyzed within the AFD-X to <1.40 Å. The map was contoured at 2σ and superimposed over a licorice representation of the protein structure.

## 5. Conclusions

To summarize, we have developed an array-type fixed-target device for X-ray crystallography (AFD-X) made from UV-curable polymers that have good optical transparency along with a high transmission factor for X-rays. We demonstrated the fabrication of these array chips in both batch and a scalable R2R fashion that has the potential of reducing the cost of manufacturing by 70-fold in terms of raw materials while accelerating the overall process 80-fold compared with traditional batch processes. The AFD-X was designed to be compatible with the SSRL crystallization plate, robot and humidity stream to allow for high-throughput as well as remote data collection at room temperature. Finally, we demonstrated the usability of the chip using three model proteins – *Gallus gallus* lysozyme, *Thaumatococcus daniellii* thaumatin and *Tritirachium album* proteinase K.

Our AFD-X platform exhibits significant potential for use in serial crystallography and/or multi-crystal data collection due to its compatibility with automation. While useful to facilitate fast and potentially remote data collection at synchrotrons, such automated sample handling is crucial for XFEL experiments where the high speed of data acquisition and the need to refresh sample is further compounded by the limited amount of beam time available. Furthermore, the optical transparency of AFD-X has the potential to enable concurrent spectroscopy and light-triggered time-resolved studies within this device. Importantly, by simplifying and streamlining the data collection process, our platform also has the potential to democratize room-temperature data collection, making this powerful approach more accessible to a broader range of researchers.

## Supporting information

PhotoMask

## 6. Acknowledgments

We would like to acknowledge Diwakaran Rathinam Palaniswamy and Bakthavachalam Kannadasan for helpful discussions, Barbara Stewart, Sondre Brandso, Dr. Jeffrey Morse and Dr. Vincent Einck for their help in roll-to-roll fabrication. We would also like to thank Jan Ilvasky at APS, Argonne National Laboratory for their help in updating Nika to integrate the diffraction images. All devices were fabricated at the University of Massachusetts Amherst Nanofabrication Cleanroom and Roll-to-Roll facility with support from the Institute for Applied Life Sciences.

This research was partially supported by a fellowship from the University of Massachusetts as part of the Chemistry-Biology Interface Training Program (National Research Service Award T32 GM139789). This investigation was also supported by a fellowship from PPG Corporation and the National Institutes of Health NIGMS R01GM149746. Use of the Stanford Synchrotron Radiation Lightsource, SLAC National Accelerator Laboratory, is supported by the U.S. Department of Energy, Office of Science, Office of Basic Energy Sciences under Contract No. DE-AC02-76SF00515. The SSRL Structural Molecular Biology Program is supported by the DOE Office of Biological and Environmental Research, and by the National Institutes of Health, National Institute of General Medical Sciences (including P41GM103393). The contents of this publication are solely the responsibility of the authors and do not necessarily represent the official views of NIGMS or NIH.

