## Supplementary Text for "Scalable Fabrication of an Array-Type Fixed-Target Device for Automated Room Temperature X-ray Protein Crystallography"

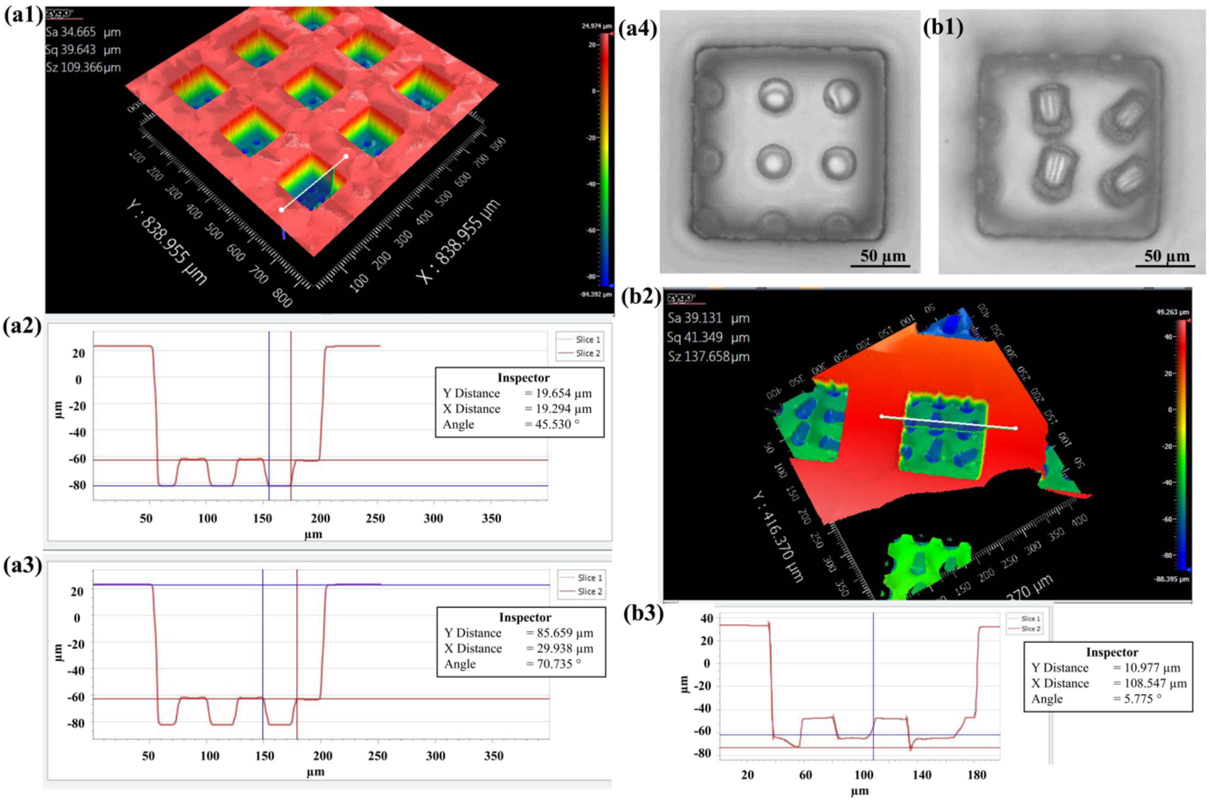


**Figure S1. (a1)** 3-D profilometry image of the NOA grids fabricated by R2R processing **(a2-a3)** 2-D cross sectional section of a single grid. **(a4)** Optical micrograph of a single grid fabricated using no added pressure. **(b1)** Optical micrograph of polymer grid fabricated using higher pressure showing deformations. **(b2-b3)** 3-D and 2-D profilometry image of the deformed NOA chip.

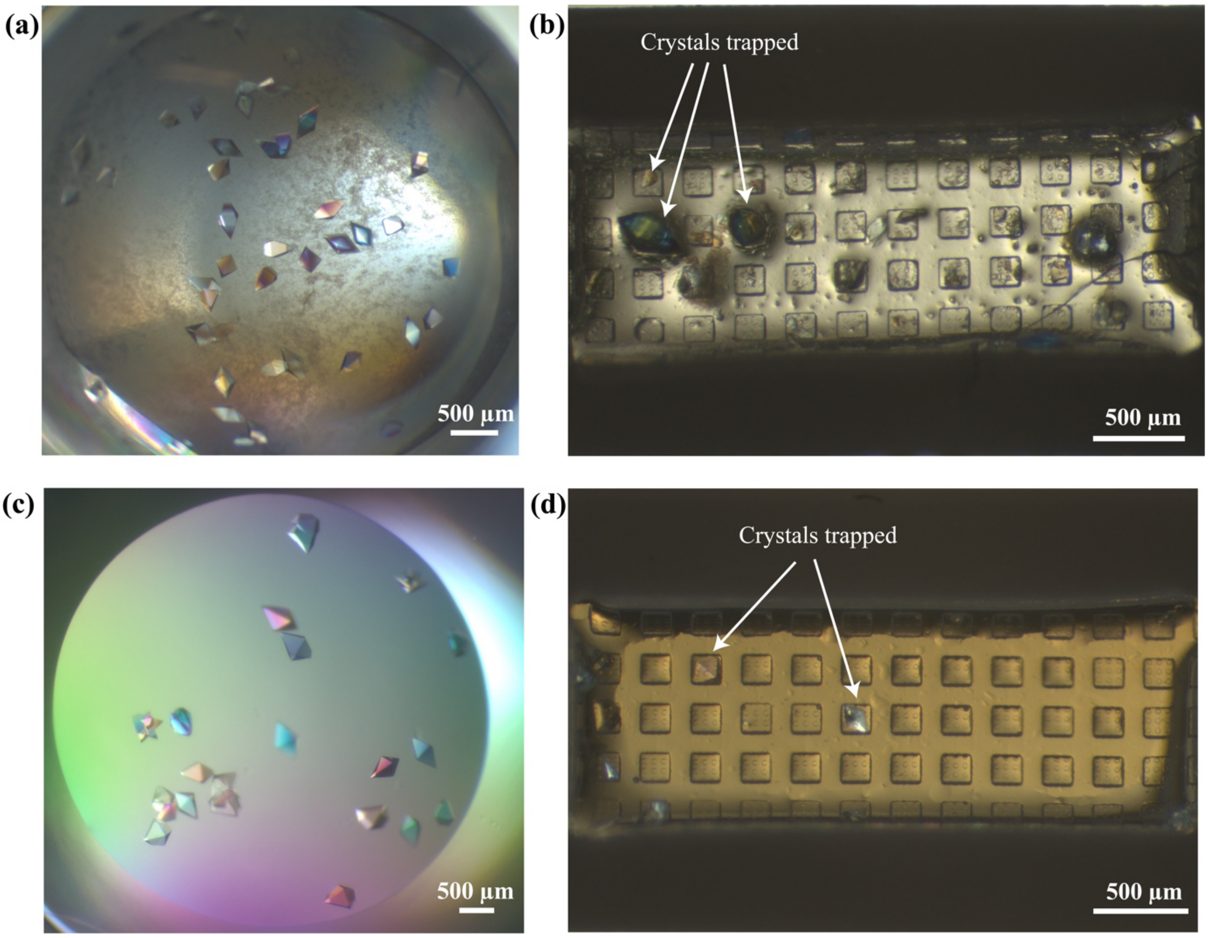


**Figure S2.** Polarized light microscopy images of **(a)** thaumatin crystals grown via sitting drop and **(b)** transferred into the AFD-X chip. **(c)** Proteinase K crystals grown via sitting drop and **(d)** transferred to the AFD-X chip.

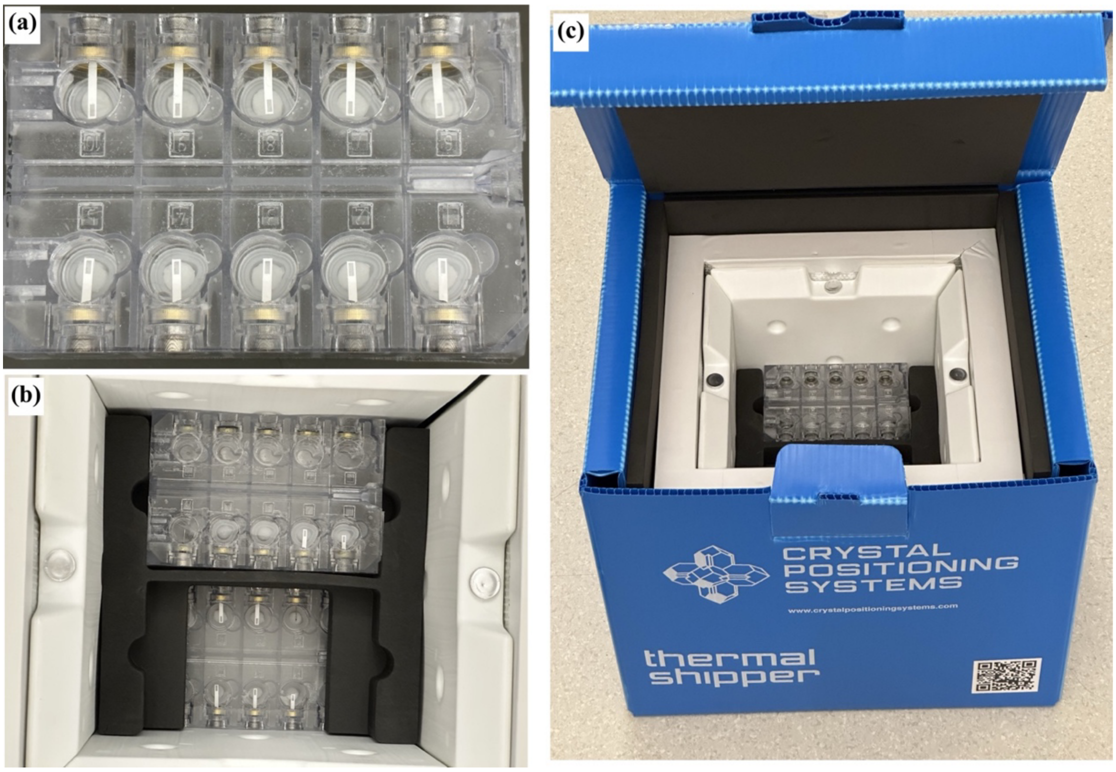


**Figure S3. (a)** Photograph of AFD-X chips loaded with protein crystals inside the SSRL crystallization plate for safe transport. **(b)** Multiple loaded plates stacked into the thermal shipper box. **(c)** Overview image of the thermal shipper box containing the SSRL plates.

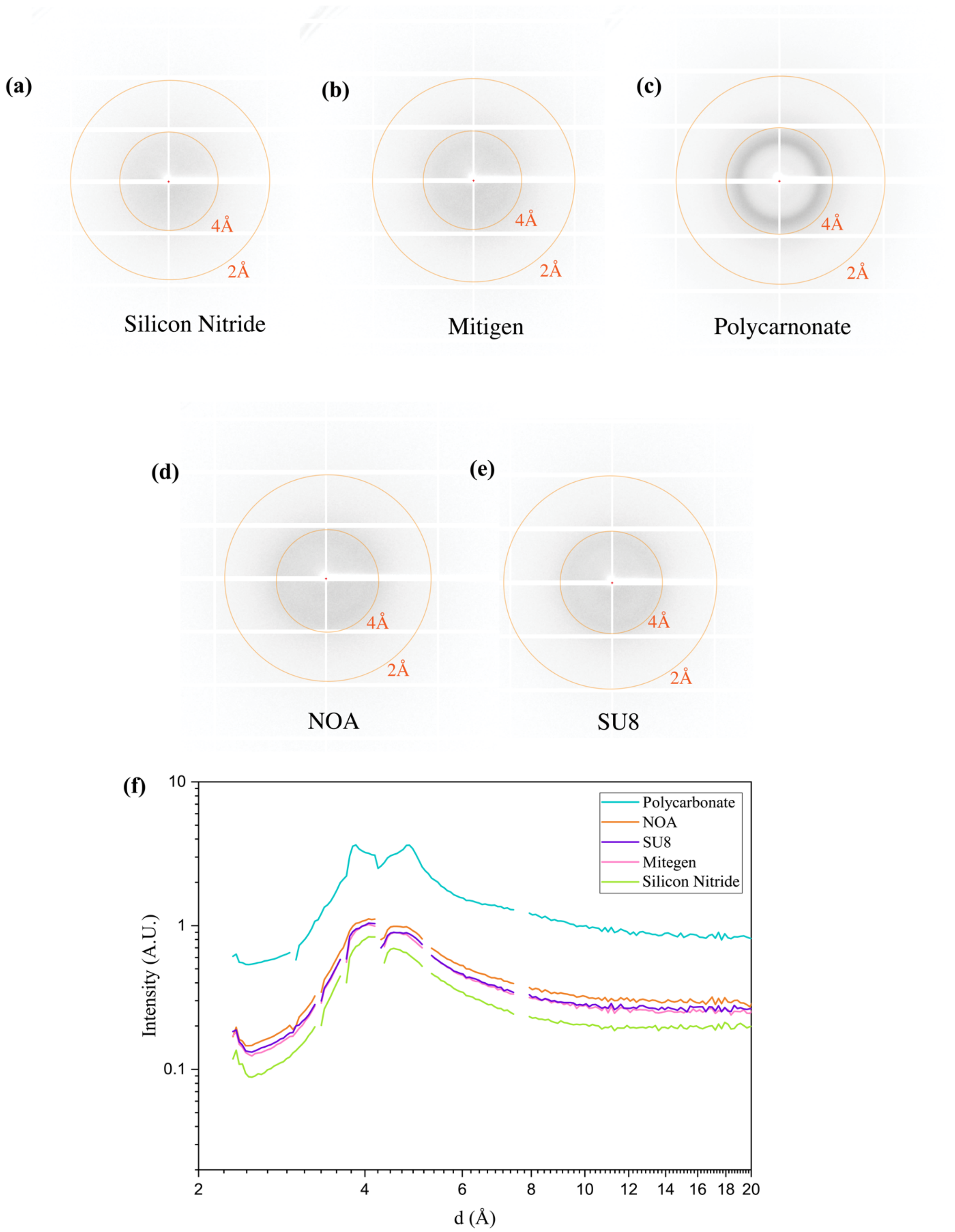


**Figure S4.** The 2-D diffraction signal comparing the commercially available fixed-target devices **(a)** made with 50 nm-thick silicon nitride,[1] **(b)** Mitigen Micromeshes, **(c)** ~200 µm-thick polycarbonate chips,[2] and AFD-X developed using **(d)** 2 µm SU-8 and **(e)** 20 µm NOA. **(f)** Graph of the 1-D integrated X-ray intensity profiles comparing the relative strength of the observed diffraction signal from the materials shown in **(a)**-**(e)**. Data were collected at SSRL beamline 12-1.

### S1. X-ray Crystallography and Crystal Isomorphism

We analyzed the statistical variation of the unit cell parameters for lysozyme, thaumatin and proteinase K crystals. After indexing and geometrical refinement of each crystal, the analysis was performed using methods described by Q. Liu *et al.*[3,4] A standard Euclidean distance, $\Delta_{j,k}$ was used to calculate the variation in crystal unit cell parameter. The Euclidean distance is a normalized distance using the unit cell parameter a, b, c, α, β, and γ of crystal *j* and *k* across *N* different crystals. It is normalized using the variance of these parameters over the population.

$$\Delta_{j,k} = \left[\frac{1}{\sigma^2}\sum_{u=\mathrm{a,b,c,}\alpha,\beta,\gamma}(u_j - u_k)^2\right]^{\frac{1}{2}} \quad \text{(S1)}$$

$$\sigma^2 = \left[\frac{1}{N}\sum_{k=1}^{N}(u_k - \bar{u}_k)^2\right] \quad \text{(S2)}$$

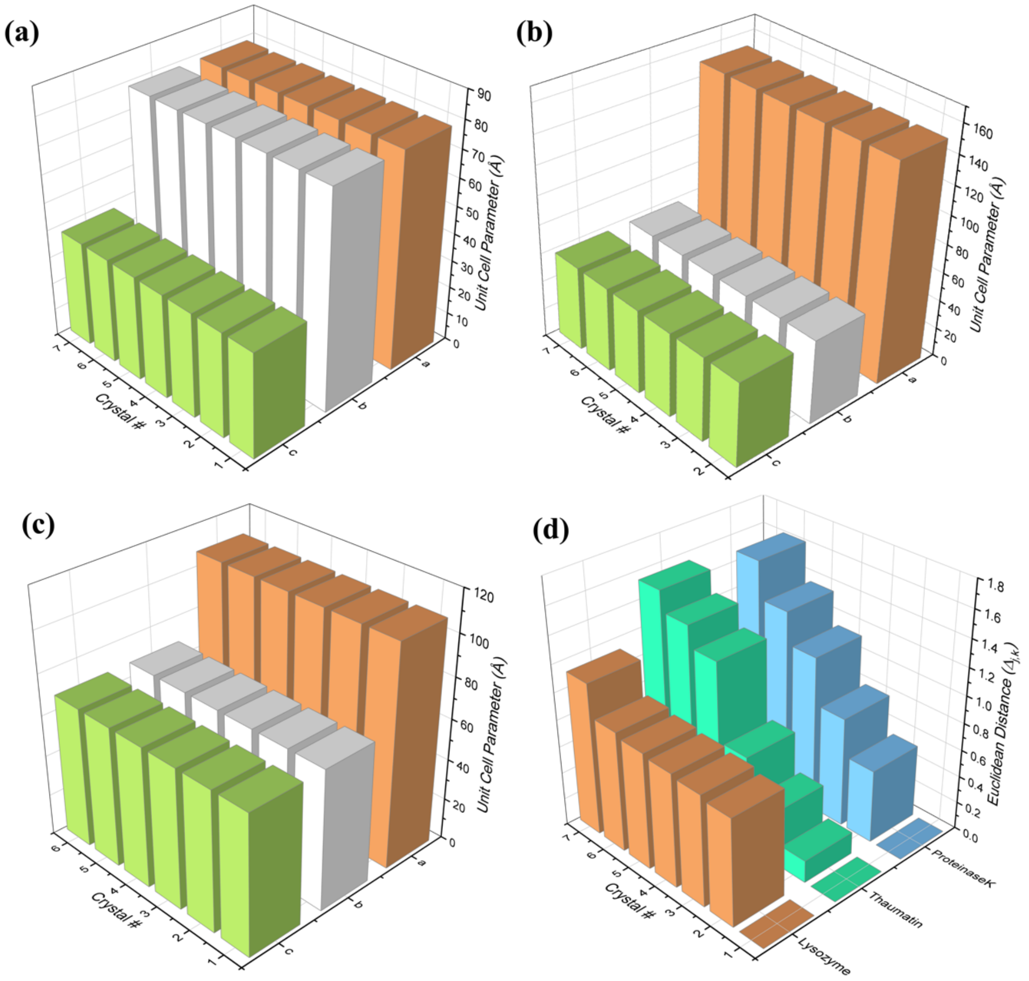


**Figure S5. (a)-(c)** Plots of the variation in the corresponding unit cell parameter across the 7 crystals of lysozyme, thaumatin and proteinase K analyzed, and **(d)** a plot of the standard Euclidian distance $\Delta_{j,k}$ vs. crystal, showing the variation in the unit cell parameters of the three different protein crystals.
